# Spatial accumulation of salicylic acid is regulated by RBOHD in potato immunity against viruses

**DOI:** 10.1101/2020.01.06.889998

**Authors:** Tjaša Lukan, Maruša Pompe-Novak, Špela Baebler, Magda Tušek-Žnidarič, Aleš Kladnik, Maja Križnik, Andrej Blejec, Maja Zagorščak, Katja Stare, Barbara Dušak, Anna Coll, Stephan Pollmann, Karolina Morgiewicz, Jacek Hennig, Kristina Gruden

## Abstract

While activation of resistance (R) proteins has been intensively studied, the downstream signaling mechanisms leading to restriction of pathogen remain mostly unknown. We studied the immunity network response conditioned by the potato *Ny-1* gene against potato virus Y. We analyzed the processes in the cell death zone and surrounding tissue on the biochemical and gene expression levels to reveal spatiotemporal regulation of immune response. We show that the transcriptional response in the cell death zone and surrounding tissue is dependent on salicylic acid (SA). For some genes, spatiotemporal regulation is completely lost in SA-deficient line, while the others show different response, indicating multiple connections between hormonal signaling modules. The induction of NADPH oxidase RBOHD expression occurs specifically on the lesion border during resistance response. In plants with silenced RBOHD, the functionality of resistance response is perturbed and virus spread is not arrested at the site of infection. RBOHD is required for spatial accumulation of SA, and conversely RBOHD is under transcriptional regulation of SA signaling. Using spatially resolved RNA-Seq, we also identified spatial regulation of an UDP-glucosyltransferase, another component in feedback activation of SA biosynthesis, thus deciphering a novel aspect of resistance signaling.

## Introduction

Plants have evolved sophisticated mechanisms to perceive pathogen attack and effectively respond, either passively or by multi-layered actively-induced immune system. The first, most general layer, pathogen-associated molecular-pattern-triggered immunity (PTI), is based on the recognition of pathogen surface components, common for a number of pathogens, by the plant pattern recognition receptors. The more specific layers of immunity are mediated by intracellular resistance (R) proteins (Jones and Dangl, 2006). R proteins confer recognition of pathogen-derived effectors and initiate effector-triggered immunity (ETI). One of the manifestations of a successful immune response is the hypersensitive response (HR)-conferred resistance, where restriction of pathogens to the infection site is associated with a form of localized programmed cell death (PCD) (Künstler et al., 2016). HR-conferred resistance is preceded by a series of biochemical and cellular signals. One of the earliest hallmarks of HR is the rapid and intense production of reactive oxygen species (ROS) (Balint-Kurti Peter, 2019). Salicylic acid (SA) is required for the restriction of pathogens during HR in various pathosystems including viruses (Mur et al., 2008; Künstler et al., 2016; Calil and Fontes, 2017) such as tobacco-tobacco mosaic virus (TMV) (Chivasa et al., 1997; Chivasa and Carr, 1998) and potato-potato virus Y (PVY) (Baebler et al., 2014; Lukan et al., 2018). The effectiveness of downstream events in ETI is regulated also by jasmonic acid (JA) and ethylene (ET). Other hormones were also shown to play important roles in plant immunity (Verma et al., 2016). Activation of immune signaling network results in the induced expression of actuators of defense, such as pathogenesis-related protein 1 (PR1) and beta-glucosidases, yet their function is not fully understood (Breen et al., 2017). HR associated PCD was shown to restrict pathogen spread in some biotrophic pathosystems (Dickman and Figueiredo, 2013). It has been however shown that it is not required for resistance in several viral pathosystems (reviewed in Künstler et al., 2016), including potato-PVY interaction (Lukan et al., 2018).

Precise temporal and spatial coordination of induced signaling pathways is required to successfully restrict the pathogen with minimal damage to the host (reviewed in Künstler et al. 2016). The concentric spread, typical of many viruses, results in foci containing cells at different stages of infection (Yang et al., 2007; Rupar et al., 2015). The immune response signal is however transferred to the surrounding tissue, resulting in a gradient of response in surrounding cells (Dorey et al., 1997; Havelda and Maule, 2000; Maule et al., 2002; Yang et al., 2007).

PVY, a member of *Potyvirus* genus from family *Potyviridae*, is the most harmful virus of cultivated potatoes (Karasev and Gray, 2013) and among top 10 most economically important viruses infecting plants (Scholthof et al., 2011). The most studied type of resistance to PVY is HR-conferred resistance (Karasev and Gray, 2013). In potato cv. Rywal, HR is initiated by *Ny-1* gene and is manifested as the formation of necrotic lesions on inoculated leaves (Szajko et al., 2008). We have shown previously that the transcriptional dynamics of genes known to participate in the immune response is crucial for the efficient resistance response and that SA is the key component in the orchestration of these events (Baebler et al., 2014).

To dissect the mechanisms acting downstream of Ny-1 R protein activation in resistance response, we analyzed the processes in the cell death zone and surrounding tissue in spatiotemporal manner on the biochemical and gene expression levels. To evaluate the position of SA in the signaling cascade, we in parallel analyzed the responses in SA-depleted NahG plants. We show that SA regulates expression of several immune-related genes in different ways. We also show that expression of few genes is induced only in the border region of viral foci, the most pronounced is the one of *RBOHD* gene. Indeed, silencing of *RBOHD* breaks the resistance, allowing systemic viral spread, which confirms that RBOHD is a regulatory hub in potato immune response. SA is required for efficient regulation of RBOHD, and conversely, RBOHD is required for spatial regulation of SA accumulation. RNA-Seq analysis of lesion and the adjacent tissue revealed that UDP-glucosyltransferase UGT76D1, an enzyme participating in feedback activation loop of SA biosynthesis, is under the regulation of RBOHD. As SA is modulating the response of genes participating in JA and ET signaling in a diverse spatiotemporal manner, we conclude that the efficient resistance response is a result of tightly interconnected hormonal network.

## Results

### Transcriptional response of immune signaling-related genes is diversely spatially regulated

Previously, we performed a non-targeted transcriptional analysis of inoculated homogenized leaves at 1, 3 and 6 dpi (Baebler et al. 2014). While conducting several independent experiments, we observed that responses can vary between plants, also related to the number of lesions formed on the leaves. We thus hypothesized that more detailed spatial analysis of responses would provide better insights into the processes involved in resistance response downstream Ny-1 activation. The symptoms development in HR interaction of potato inoculated with PVY^N-Wilga^ (cv. Rywal, hereafter NT) and its transgenic counterpart depleted in accumulation of SA (NahG-Rywal, hereafter NahG) was as previously described (Baebler et al., 2014; Lukan et al., 2018). While resistance was successful in NT and the virus was restricted to inoculated leaves, depletion of SA rendered NahG plants susceptible allowing the viral spread throughout the plant accompanied by strong symptom development (Figure S1).

We developed the protocol for precise sampling of tissue sections surrounding the lesion at different stages of lesion development (Figure 1a) suitable for transcriptomics and hormonomics analysis. As the amounts of RNA obtained from individual small tissue sections (caa. 400 cells) (Figure S2) are limited, we first selected 23 genes for our analysis according to their biological function and responsiveness in our microarray dataset (Baebler et al. 2014; Table S1) participating in redox potential homeostasis (*CAT1*, *PRX28*, *RBOHD*, *RBOHA*, *RBOHC*, *TRXO*, *TRXH*), cell death (*MC3*, *LSD1*), RNA silencing (*AGO2*, *SAHH*), MAP kinase cascade (*WIPK*, *MKP1*), SA (*TGA2*), JA (*13-LOX*, *9-LOX*, *ACX3*), ET (*ERF1*) or auxin (*ARF2*) signaling, being actuators of defense (*BGLU*, *PR1B*) or markers for primary metabolism state (*GBSSI*) (Figure 1c). In parallel to gene expression of those genes, the relative amount of PVY RNA was measured (Data S1, S2).

**Figure 1:**
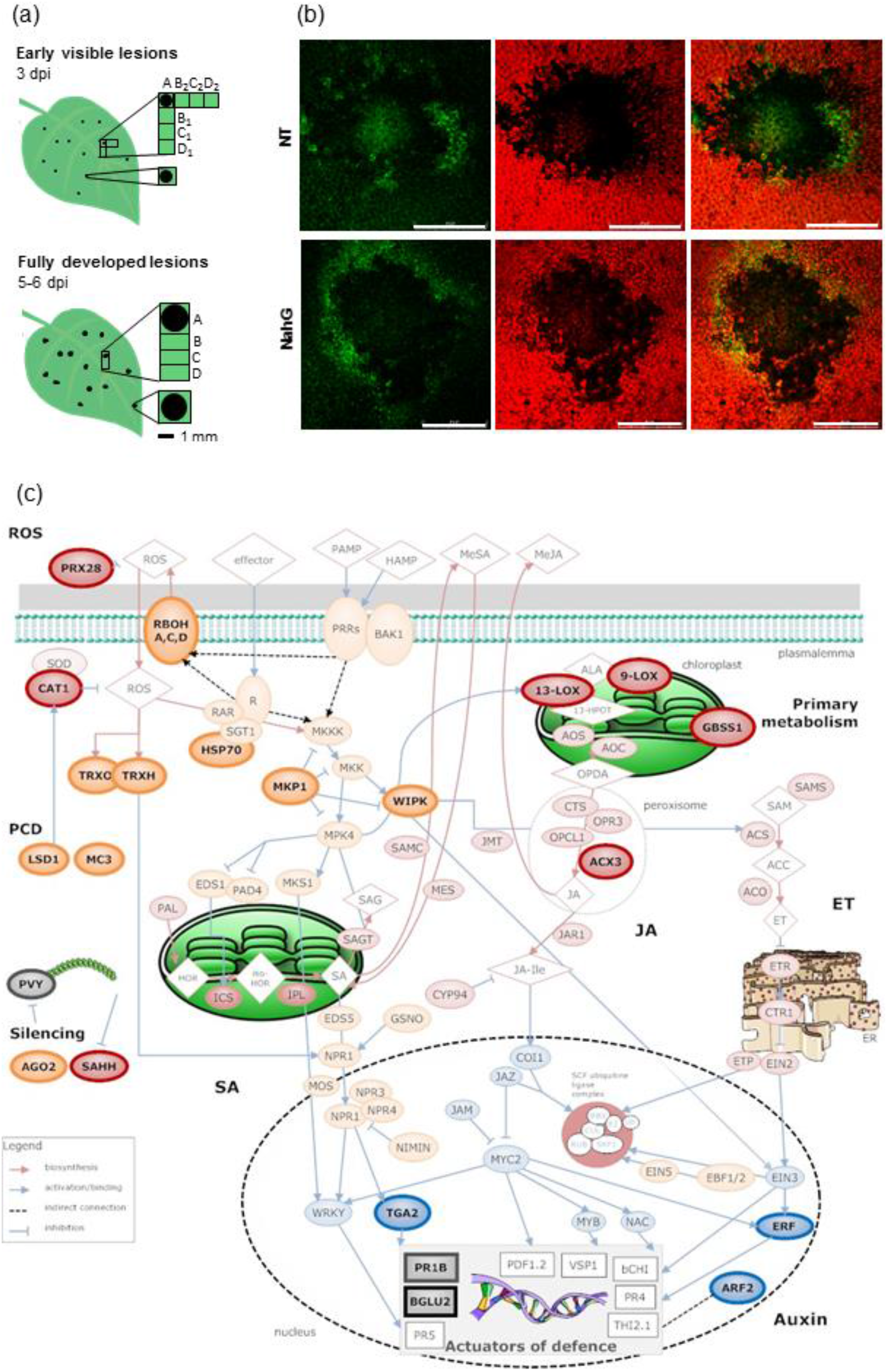
Experimental setup. (a) Sections containing lesion (section A) and surrounding tissues (sections B, C and D, consecutive 1 mm strips distal to the section A) were sampled at two different time points. Tissue surrounding early visible lesions was sampled in two perpendicular directions, here marked with 1 and 2, respectively. In mock-inoculated plants, 4 adjacent tissue sections of the same sizes as in inoculated leaves were excised from different plants. Scale bar: 1 mm. (b) PVY^N^-GFP accumulation around fully developed lesions in non-transgenic (NT, top) or in SA-depleted (NahG, bottom) plants. From left to right: PVY^N^-GFP accumulation (green), chlorophyll fluorescence (red), overlay of chlorophyll fluorescence and PVY^N^-GFP accumulation. Scale bar: 2 mm. (c) Plant immune signaling-related genes selected for the initial transcriptomic spatiotemporal response analysis. Potato virus Y (PVY) accumulation and expression of genes involved in reactive oxygen species (ROS), MAP kinase signaling, programmed cell death (PCD), salicylic acid (SA), jasmonic acid (JA), ethylene (ET), auxin and primary metabolism were analyzed (bold border). Signaling proteins are presented as orange, enzymes as red and transcription factors as blue ovals, respectively; metabolites are presented as diamonds and genes as squares. 13-LOX: 13-lipoxygenase, 9-LOX: 9-lipoxygenase, ACX3: acyl-CoA oxidase, AGO2: argonaute 2, ARF2: auxin response factor 2, BGLU2: 1,3-beta-glucosidase, CAT1: catalase 1, ERF1: potato ethylene responsive transcription factor 1a, GBSS1: granule-bound starch synthase 1, HSP70: heat shock protein 70, LSD1: zinc finger protein LSD1, MC3: metacaspase 3, MKP1: mitogen-activated protein kinase phosphatase 1, PR1B: pathogenesis-related protein 1b, PRX28: peroxidase 28, RBOHA: potato respiratory burst oxidase homologue A, RBOHC: potato respiratory burst oxidase homologue C, RBOHD: potato respiratory burst oxidase homologue D, SAHH: adenosylhomocysteinase S-adenosyl-L-homocysteine hydrolase, TGA2: bZIP transcription factor family protein, TRXH: thioredoxin H, TRXO: thioredoxin O, WIPK: potato wound inducible protein kinase. See Table S4 for gene IDs and primer information.

First, we analyzed fully developed lesions following the infection with PVY^N-Wilga^ (Figure 1a). While there was most of viral RNA in the centre of the lesion in both potato genotypes, the expression pattern of genes changed with distance from the centre of the lesion and in some cases, the response was different between the two genotypes (Figure 2). The response did however not differ if the plant was inoculated with different viral strains (PVY^NTN^, PVY^N^-GFP, Figure S3). To simplify the visualization and interpretation of spatial response comparisons among different genes in the two genotypes, we designed coefficient plots, condensing the information of spatial response to pairs of quadratic and linear coefficient values from the polynomial model fit (Figure 2, bottom position). The expression of *HSP70* (see Figure 1c for full gene names) encoding a protein involved in the stability of R proteins (Kadota and Shirasu, 2012), is elevated in the centre of viral foci (section A) compared to the adjacent sections (sections B, C and D) only in non-transgenic (NT) plants (Figure 2). Both, *BGLU2* and *PR1B* (Figure 2) are also under the regulation of SA signaling pathway. While they indeed show strong upregulation close to the centre of viral infection (more than 10-fold compared to the distal tissue, D section) during resistance response, their response quite differs in SA-depleted system. While the spatial response of *BGLU2* in NahG plants is completely lost, the peak of *PR1B* response is occurring further away from the foci of viral infection (peak in section C, Figure 2). The regulation of *ERF1*, involved in ET signaling, resembles the one of *PR1B* in both genotypes. The starch synthase *GBSS1*, an indicator of primary metabolism state, shows that the metabolism is downregulated in the centre of the lesion and its proximity, while in section D, its expression status is closer to the one in mock-inoculated leaf tissue (Figure 2, denoted with empty circles at the right end of x-axis). This is however not true in NahG plants where the expression of *GBSS1* is downregulated even further away from the lesion (Figure 2).

**Figure 2:**
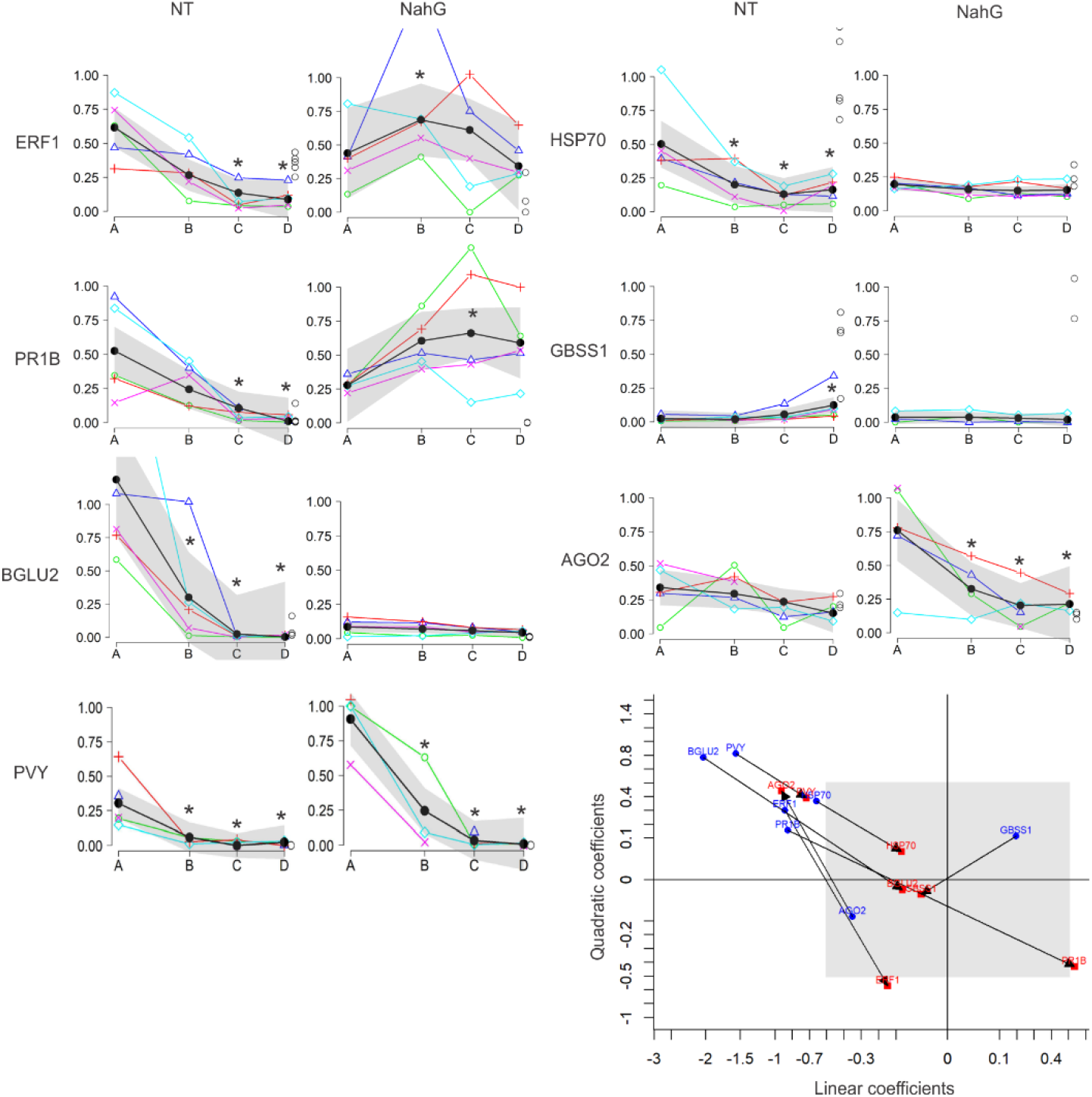
The lack of SA leads to diverse spatial transcriptional regulation of immune-related genes. Transcriptional profiles of the selected genes were monitored in non-transgenic (NT) and SA-depleted (NahG) plants following inoculation with PVY^N-Wilga^ at the stage of fully developed lesions (top position). Tissue sections, marked as positions A, B, C, D, are shown on the x-axis and values of gene transcriptional response standardized to 97^th^ quantile on the y-axis. Expression values in mock-inoculated tissue sections are shown as empty circles on the right end of x-axis. Asterisk denotes statistical significance (p-value < 0.05) of gene expression at position B, C and D compared to that at position A. Spatial profile models are shown as thick black lines with 95% confidence interval bands in grey. Standardized gene expression values within individual lesions are presented with colored symbols connected by a line. To simplify the visualization and interpretation of spatial response comparisons among different genes within different genotypes, we designed coefficient plots (plotted pairs of quadratic and linear coefficient values from the polynomial model fit, bottom right position). Value pairs for NT are shown as blue circles, value pairs for NahG as red squares. Grey box defined within [−0.5,0.5] interval boundaries indicates an area of lower relevance regarding the difference in gene expression response. Full gene names are given in the legend of Figure 1c.

We also show that expression of genes involved in JA biosynthesis, maintenance of redox potential and the ones involved in cell death is spatiotemporally regulated, which is presented in more detail in next subchapters. The rest of the analyzed genes showed either no spatial regulation at all or they were responding in an asynchronous manner between the biological replicates (Data S1, Data S3), meaning that most probably their regulation is even more fine-tuned and more detailed analysis (e.g. with single cell resolution) would be required.

Next, we wanted to implement our approach in the earlier stages of viral infection. As the first robust time point for spatial analysis of responses, we identified the early visible lesions, these are lesions visible only when leaves are transilluminated and are of 0.5-1 mm in diameter (Figure 1a). Indeed, when comparing the response profiles of different lesions and profiles within the same lesion in perpendicular directions, the results were consistent and enabled reliable biological interpretation (Figure S4, Data S2, Data S4). For example, the JA signaling module is not uniformly spatiotemporally regulated. *13-LOX* gene, encoding the 13-lipoxygenase starting the oxylipin branch leading to JA biosynthesis, is induced in the centre of viral foci at the stage of fully developed lesions (Figure S5). The response is, in agreement with the hypothesis that JA and SA are antagonistic, enhanced in NahG plants. The expression pattern of *ACX3* gene, involved in later steps of JA biosynthesis (Figure 1c), is regulated similarly, except for the diminished regulation at the stage of fully developed lesions of NahG plants (Figure S5). Altogether, we show that spatiotemporal transcriptional response of immune related genes is not homogenous within one signaling pathway indicating tight interconnectedness between different components of immunity.

### Expression of genes involved in redox state maintenance is tightly regulated across the studied spatiotemporal scale

We next focused our research on the genes involved in generation and quenching of ROS as redox potential changes are one of the first responses to pathogen recognition both in PTI and ETI (Figure 1c). We studied three *RBOH* genes, involved in generation of apoplastic ROS and leading to redox changes in the cytoplasm. On the quenching side, we analyzed gene expression of CAT1, located in the peroxisome, and apoplastic peroxidase PRX28. Additionally, we monitored the gene expression of two cytoplasmic redox potential sensor proteins that were implicated in the regulation of immune signaling, thioredoxins H and O (TRXH and TRXO). The three investigated *RBOH* genes have each distinct spatiotemporal regulation. *RBOHC* is not regulated in any of the studied genotypes (Figure 3). *RBOHA* is strongly induced in the centre of viral foci already in early stage of lesion development, while this activation is attenuated in NahG plants. *RBOHD*, however, responds in section A in early stages while later on, it forms a peak of expression close to the border of the lesion itself in the majority of lesions (Figure 3). *RBOHD* is also under regulation of SA as this gene is expressed close to the limit of quantification in NahG plants. *PRX28* is induced earlier in NahG plants, and at the stage of fully developed lesion its expression peaks in section A in NT and in section B in NahG plants. *CAT1* is induced only in NahG plants and is induced even further away from the centre of viral foci compared to *PRX28*. At the stage of early lesions, it peaks in section C and at the stage of fully developed lesions even in section D. The expression of *TRXO* was below the limit of quantification, therefore, it was not possible to follow its response. Interestingly, however, *TRXH* responds strongly and early only in NT plants. Tight spatiotemporal regulation of ROS related genes is thus important for successful viral arrest.

**Figure 3:**
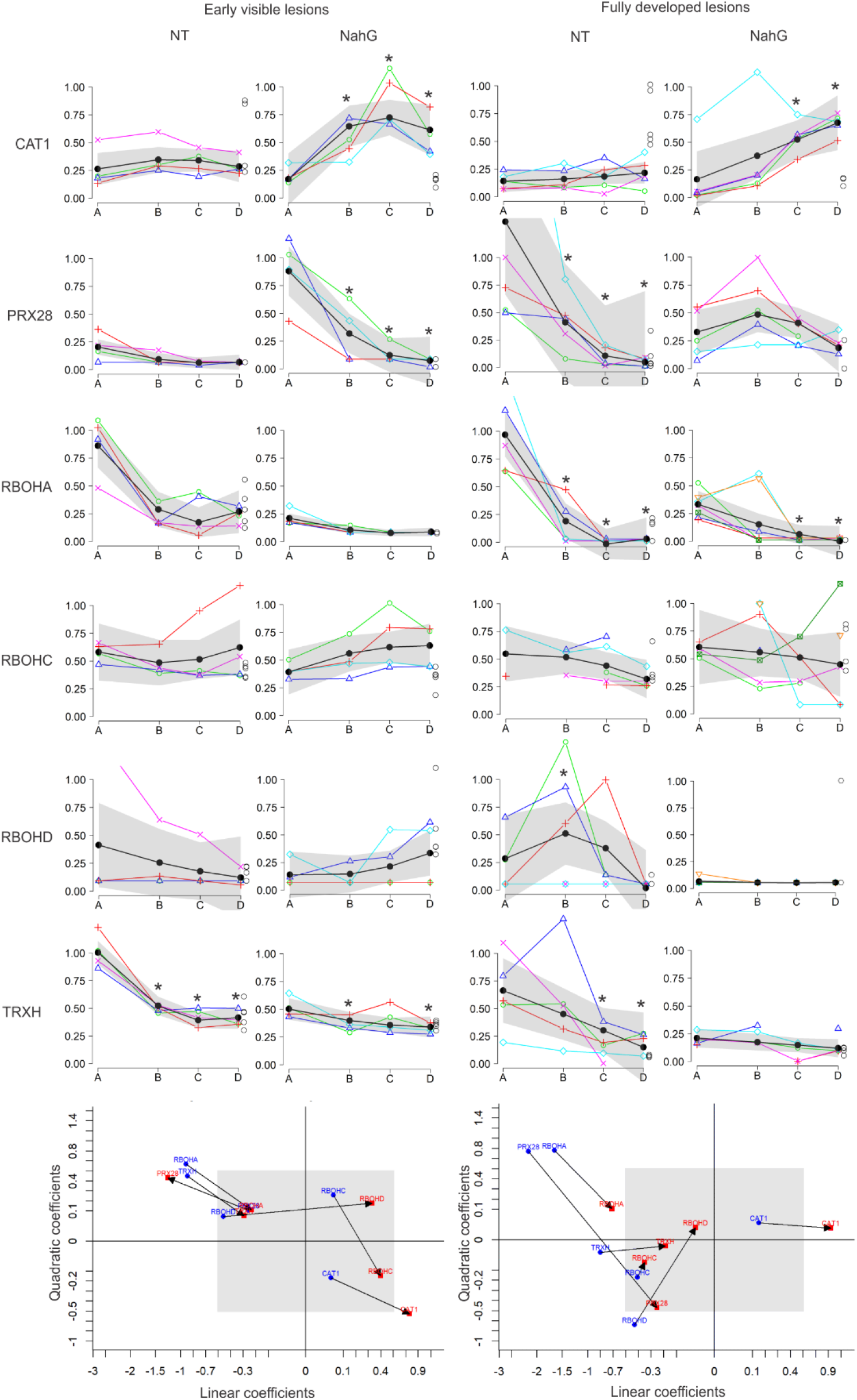
Induction of expression of *RBOHD* is limited to the cells immedately surrounding the infected tissue. Spatial expression profiles of selected genes (top position) and coefficient plots (bottom position) for non-transgenic (NT) and SA-depleted (NahG) plants after inoculation with PVY at the stage of early visible (3 dpi, left, averaged profile of the two perpendicular directions) and fully developed lesions (5-6 dpi, right). See Figure 2 legend for the full description of the graphs and Figure 1c for full gene names.

We further wanted to inspect how changes in transcriptional activity of ROS-related genes are reflected in accumulation of H_2_O_2_. We followed H_2_O_2_ production in relation to lesion size in several time points following virus inoculation. The H_2_O_2_ accumulation area around the lesions was significantly larger in NahG plants after 5 dpi (Figure 4).

**Figure 4:**
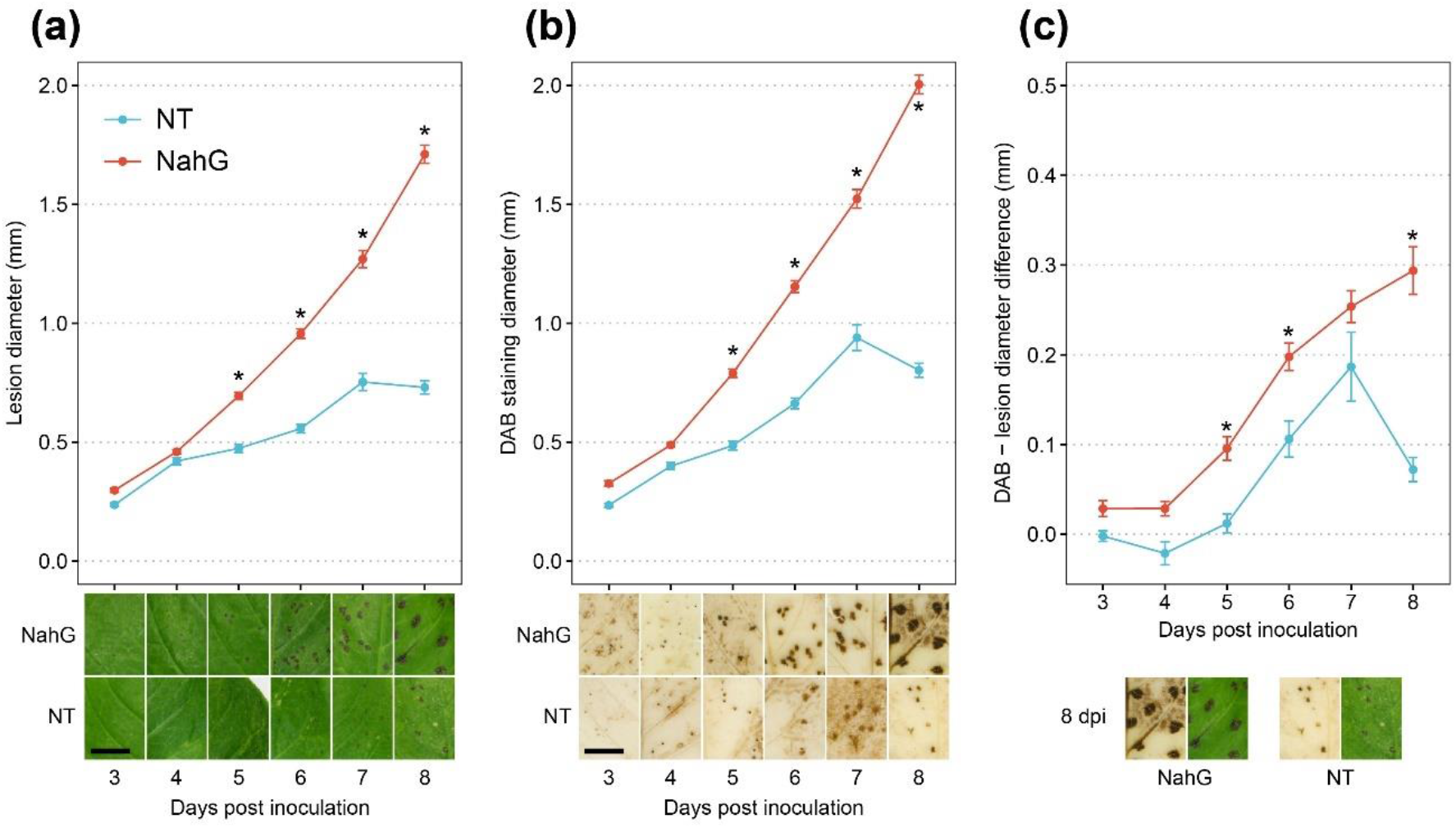
H_2_O_2_ accumulation is continuously stronger during infection with PVY in potato plants perturbed in SA signaling. (a) Lesion diameter was significantly larger in the NahG plants from 5 dpi on compared to non-transgenic (NT) ones. (b) DAB staining indicating H_2_O_2_ activity (visible as the brown precipitate) was significantly wider in NahG plants from 5 dpi on, if compared to NT plants. (c) Difference in diameter of DAB staining and the diameter of the corresponding lesion showing that in NahG plants the H_2_O_2_ positive area outside the lesion is wider in comparison to NT plants. The representative areas of leaf images and comparison of lesion and DAB staining at 8 dpi are shown under the plots (bar represents 10 mm). Asterisks represent statistically significant differences between the two genotypes at the same day (p < 0.05). Error bars represent standard error.

### Functional *RBOHD* is indispensable in resistance against PVY

*RBOHD* showed a specific spatiotemporal expression profile, peaking at the border of virus amplification zone (Figure 3). To validate the involvement of ROS signaling and specifically *RBOHD* gene in virus restriction, we constructed transgenic plants of cv. Rywal with silenced *RBOHD* gene (shRBOHD; Figure S6). We inoculated two independent transgenic lines of shRBOHD (lines 13 and 14) with PVY^N^-GFP as the least virulent PVY strain among the ones used in this study (Lacomme et al., 2017). The virus spread systemically and lesions appeared in upper, non-inoculated leaves (Figure 5a, Figure S6). Time of first lesion appearance in upper, non-inoculated leaves was comparable to the time of first lesion appearance in NahG plants, but the viral RNA amounts were lower than in NahG plants (Figure S6). We also checked if the virus can multiply to a larger extent in the inoculated leaves of shRBOHD plants. The viral amount negatively correlated (r = −0.85) with *RBOHD* expression taking into account all analyzed genotypes (Figure 5, Figure S6). Fully functional RBOHD thus contributes both to cell-to-cell and long distance movement of the virus.

**Figure 5:**
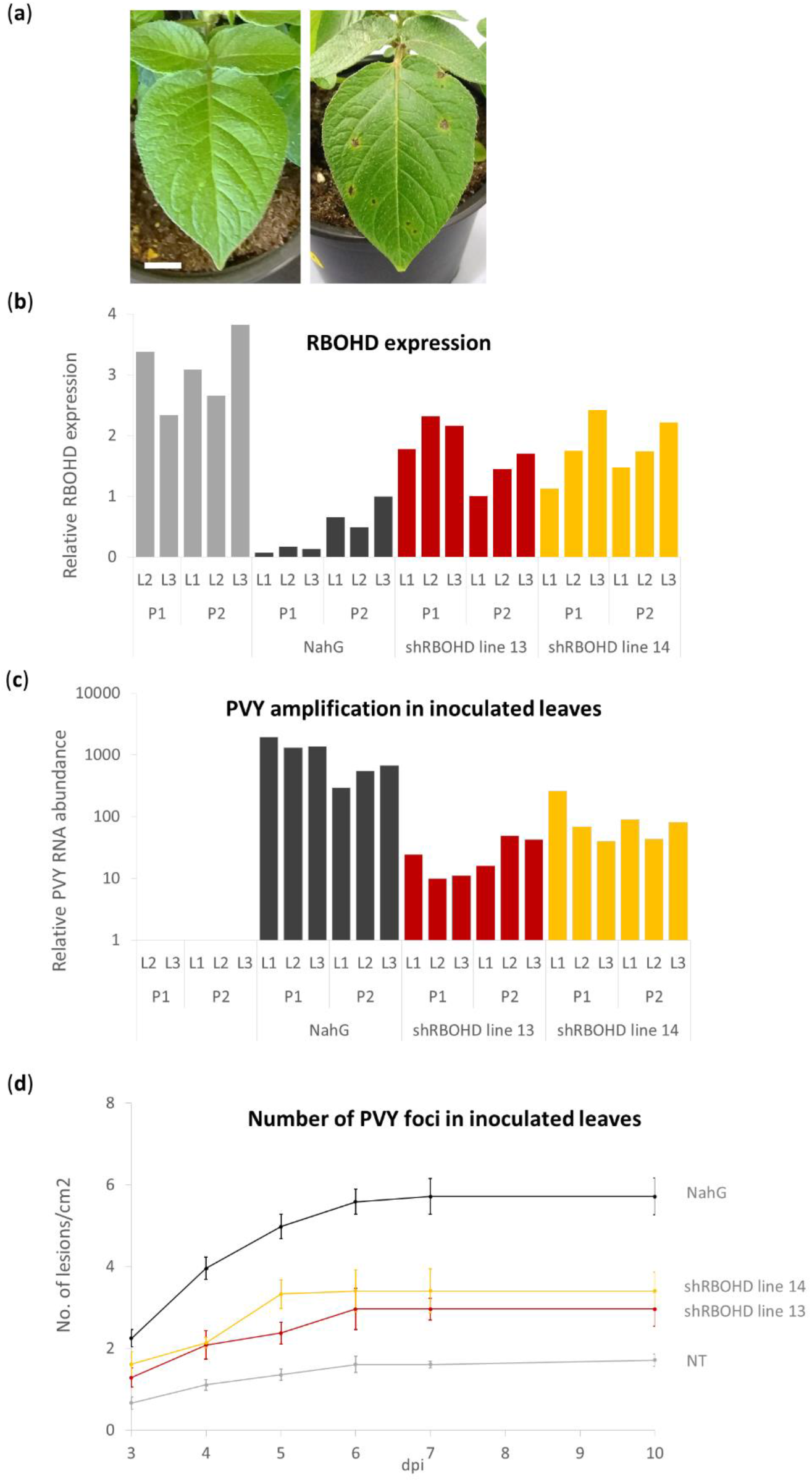
Virus efficiently multiplies in plants with silenced *RBOHD* gene. (a) Lesions on upper non-inoculated leaf of *RBOHD*-silenced plant of transgenic line 14 (right) and non-transgenic counterpart (left) 17 dpi. Scale bar is 1 cm. (b) Relative *RBOHD* expression in inoculated leaves 6 dpi. (c) Relative PVY RNA abundance (relative to the lowest detected viral amount, shown on a logarithmic scale) 6 dpi in NT, NahG and shRBOHD plants. P: plant number, L: leaf number (d) Number of lesions per cm^2^ as they appeared on the youngest inoculated leaves of plants (n = 6 – 10) of different genotypes at particular dpi. Error bars represent the standard error of the number of lesions per cm^2^. Individual measurements are available in the Data S5.

We additionally followed the number of lesions appearing on the inoculated leaves in the *RBOHD*-silenced transgenic lines after inoculation with PVY in comparison with NT and NahG plants. We observed an increased number of viral foci on inoculated leaves of shRBOHD plants in comparison to NT plants (Figure 5d, Figure S7). We conclude that the RBOHD also contributes to blocking of viral multiplication within the cell and thus establishment of foci of viral infection.

### RBOHD is involved in tight spatial regulation of SA accumulation during resistance response

ROS were shown to be involved both upstream and downstream of SA signaling (Herrera-Vasquez et al., 2015). To determine if RBOHD-generated ROS are involved in regulation of SA signaling in resistance response, we analyzed SA induction in shRBOHD plants. In the pool of lesions (section A), SA was induced to the same extent as in RBOHD-silenced plants (approximately 10-fold compared to mock samples; Figure 6, Table S2), while in the surrounding cells (sections B) accumulation was lower in non-transgenic plants but not in RBOHD-silenced plants. Since the concentration of SA in transgenic plants did not differ significantly from the concentration in non-transgenic plants in the lesion (Figure 6), we conclude that RBOHD-generated ROS are not involved in the induction of SA biosynthesis. However, the tight spatial regulation of SA biosynthesis was lost in shRBOHD plants. Additionally, we know that *RBOHD* gene activity is under regulation of SA as the expression of this gene is much lower in NahG plants after virus inoculation (Figure 5b, Figure S6, Data S1).

**Figure 6:**
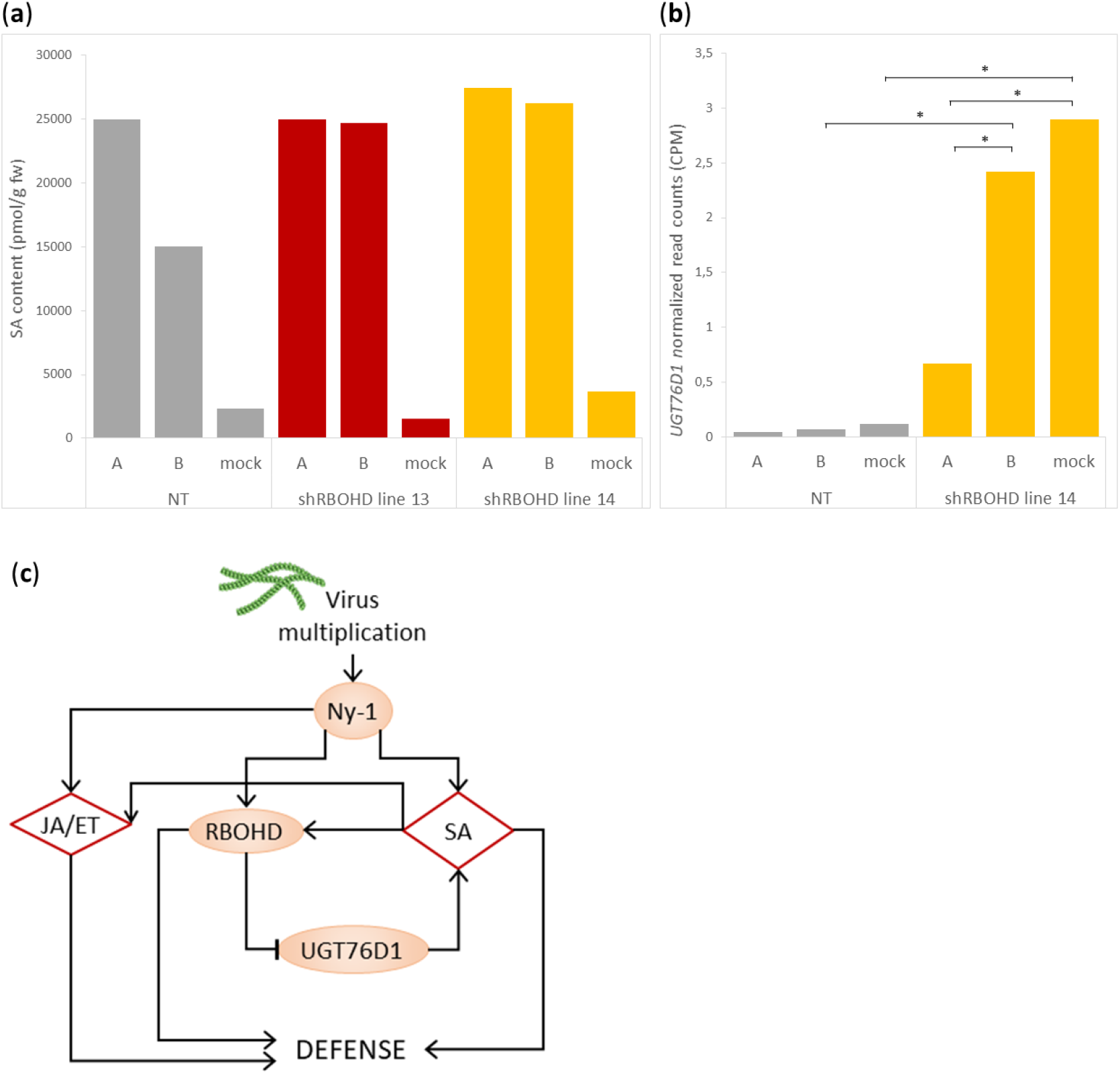
Functional *RBOHD* contributes to spatial regulation of salicylic acid accumulation in potato resistance against PVY. (a) Salicilic acid (SA) content was measured in the early stage of lesion development in non-transgenic (NT) and RBOHD-silenced (shRBOHD) plants within the lesion (section A) and in the tissue surrounding the lesion (section B) in comparison to mock-inoculated sections. Pools of 50-100 tissue sections from 6-8 plants were analyzed. Detailed data is shown in Table S2. (b) Expression of *UDP-glucosyltransferase* (*UGT76D1*) within the lesion (section A), its close surroundings (section B) and sections of mock-inoculated plants of NT and shRBOHD plants, as determined by RNA-Seq analysis. The bar graph is showing the average normalized read counts (CPM) of *UGT76D1* gene across three biological replicates. Asterisks represent statistically significant differences among comparisons (FDR adj. p-value < 0.05). (c) SA synthesis and *RBOHD* are interconnected through regulatory loops. SA regulates *RBOHD* expression and on the other hand, *RBOHD* is, through *UGT76D1*, involved in spatial distribution of SA accumulation. Signaling proteins are presented as orange ovals and metabolites as red diamonds. JA/ET (jasmonic acid/ethylene signaling), *Ny-1*: R protein.

We have further explored the mechanisms underlying spatial regulation of SA accumulation on the border of the lesion. We performed RNA-Seq analysis of the tissue within the lesion (section A) and the adjacent tissue (section B) at the time of early visible lesions for all three genotypes, NT, NahG and shRBOHD. This analysis provided insights into processes regulated by SA or RBOHD (Figure 7, Figure S8, Data S6). The majority of processes spatially differentially regulated overlap between NahG and shRBOHD plants, as *RBOHD* expression is repressed in NahG plants. We, however, also show that some of the genes and even processes/protein groups are specifically regulated only when *RBOHD* gene is silenced. Most notable regulation is regulation of MYB transcription factors (three regulated only in the absence of *RBOHD* and not in NT or NahG transgenic plants, Data S6) and regulation of protein degradation (specific regulation of several proteases as well as ubiquitin SCF and RING E3 ligases, Data S6). This shows that RBOHD has some additional roles that are independent of SA.

**Figure 7:**
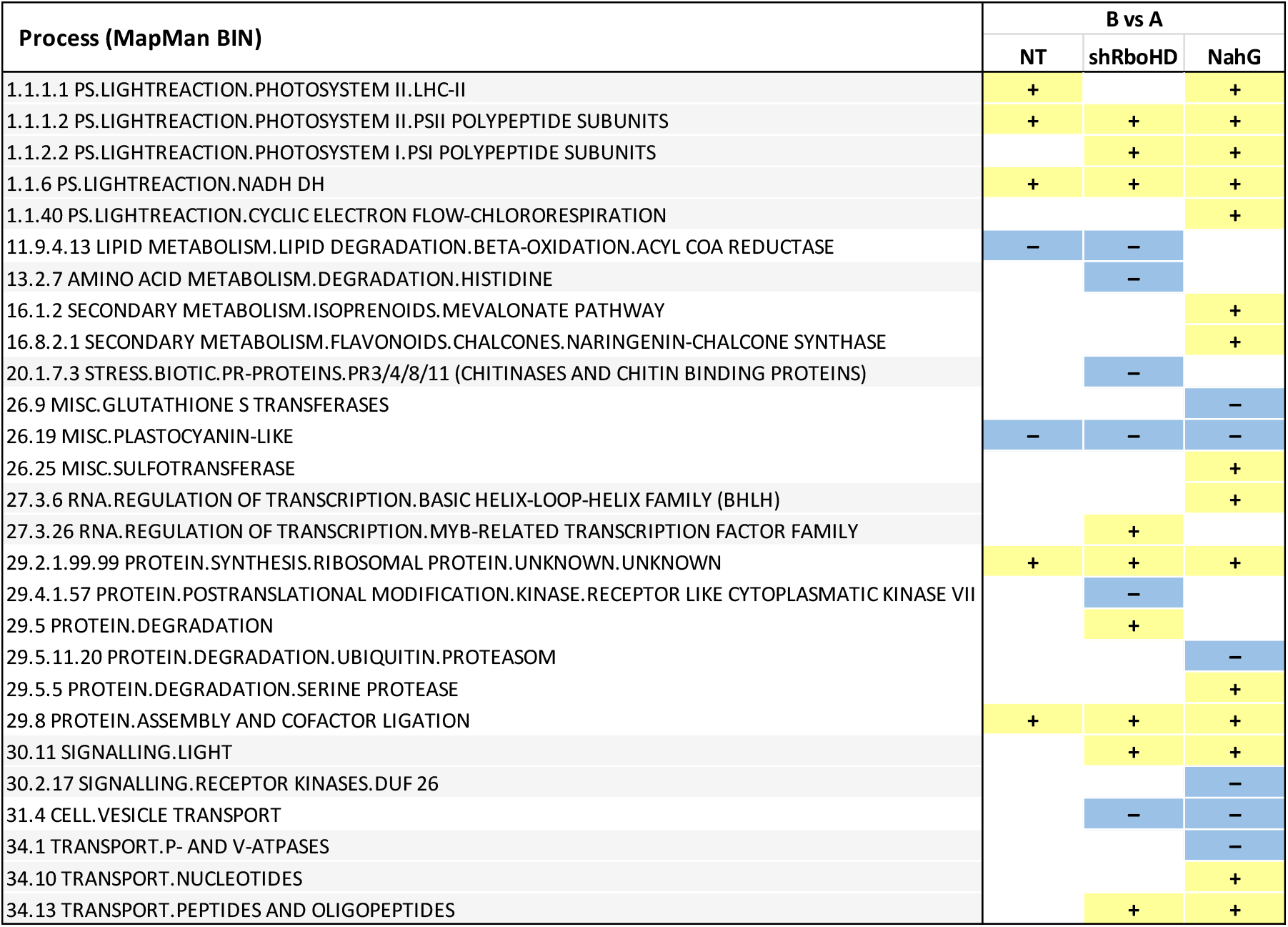
*RBOHD* contributes to spatial regulation of processes in potato resistance signaling against viruses. Differentially regulated processes in lesion (A) and tissue surrounding it (B) were compared in non-transgenic plants (NT) and in plants with perturbed SA accumulation (NahG) or reduced *RBOHD* expression (shRBOHD). Only statistically significant (false discovery rate corrected Q-value < 0.05) enriched gene sets determined by gene set enrichment analysis (GSEA) in at least one comparison are presented. “+”-induced processes, “−”-repressed processes, blanks denote that a current process was not statistically significantly enriched.

We further inspected the RNA-Seq results for any evidence of the feedback loop indicating the role of *RBOHD* in spatial regulation of SA accumulation. We have identified an *UDP-glucosyltransferase* (*UGT76D1,* Sotub10g024000) that is strongly spatially deregulated in RBOHD silenced plants and could be the one directly involved in the process (Figure 6b, Figure S9). In line with the evidence that *RBOHD* is repressed in NahG plants, similar expression pattern is observed also in these plants. UGT76D1 is in Arabidopsis implied in feedback activation loop of SA biosynthesis (Huang et al., 2018). Overexpression of this gene led to accumulation of SA. Here we similarly show that silencing of *RBOHD* results in induction of *UGT76D1* gene in section B which explains higher levels of SA detected in B section of transgenic lines (Figure 6).

## Discussion

The mechanism of activation of R proteins and their interaction with effectors was the subject of intensive research in recent years (Baggs et al., 2017; Kourelis and Van Der Hoorn, 2018; Macho and Lozano-Duran, 2019). However, to get towards sustainable resistance in the field where plant is exposed also to other environmental stressors, understanding of downstream processes leading to pathogen arrest is of importance to balance the trade-offs between growth and immunity. To date, most of the studies of those processes were performed for the model plant Arabidopsis (Piquerez et al., 2014). Although some findings can be transferred to crop species using orthology (Lee et al., 2015; Ramšak et al., 2018) this is not always the case as, for example, in recently reported redundancy of *PAD4* in *Solanaceae* (Gantner et al., 2019). Therefore it is important to perform the studies also in crop plants, such as potato.

Pathogen-infected leaf tissue comprises a heterogeneous mixture of host cells in different stages of defense response due to pathogen and defense signal movement from the primary infected cell (Maule et al., 2002). Recently, spatially-resolved transcriptome profiling in plant tissues has gained importance in unravelling complex regulatory networks (Giacomello et al., 2017; Shulse et al., 2019). The progressive and asynchronous effects of viral infection on host gene expression in relation to the spatial distribution of the virus have been investigated previously, but in compatible interactions (Aranda et al., 1996; Escaler et al., 2000; Maule et al., 2002; Yang et al., 2007). In resistance, however, the spatial distribution of a limited number of components has been studied only in model plants (Antoniw and White, 1986; Mur et al., 1997; Betsuyaku et al., 2018; Giolai et al., 2019). Our approach for analysis of small tissue sections presents a step forward in studying the resistance response against the virus and enabled identification of a novel key player and the interconnectivity of components of immune signaling.

In immunity, ROS are generated in two phases (Herrera-Vasquez et al., 2015). The first, low-amplitude and transitory phase occurs within minutes after infection. ROS generated at this stage is mostly apoplastic, tightly linked to posttranslational activation of plasma-membrane RBOH NADPH oxidases (e.g., AtRBOHD and AtRBOHF) and cell-wall peroxidases and is attributed to PTI (Torres, 2010; Shapiguzov et al., 2012; Kadota et al., 2019). The second, high-amplitude and sustained phase takes place a few hours after infection and depends on ROS generation in multiple compartments, including the apoplast, chloroplasts, mitochondria, and peroxisomes. This second phase is linked to ETI and requires transcriptional activation of *RBOH* genes (Shapiguzov et al., 2012). The role of RBOH proteins as the principal generator of ROS after pathogen attack was, using mutant and transgenic lines, mostly studied in compatible interactions with bacterial and oomycete pathogens (Allan et al., 2001; Torres et al., 2002; Yoshioka et al., 2003; Peer et al., 2011), but also in symbiotic interaction with rhizobia (Yu et al., 2018). Our spatiotemporal analysis of responses revealed that potato *RBOHD* (orthologue of *AtRBOHD*) is transcriptionally induced at the border region between virus-replicating and healthy cells around the cell death zone in resistance response (Figure 3). Several studies pointed to the essential role of ROS also in virus resistance in HR (reviewed in Hernández et al. 2015), including one of the most studied viral pathosystems, TMV-tobacco interaction (Mur et al., 1997; Liao et al., 2015). While the induction of *RBOHD* in potato-PVY interaction was shown before (Otulak-Kozieł et al., 2019), its role in resistance response was not functionally studied. We directly confirmed its essential role in potato resistance signaling, as RBOHD silenced transgenic plants were not able to arrest the virus spread (Figure 5). Besides the effect on cell to cell movement, both SA and *RBOHD* have also an effect on the efficiency of viral infection. The number of established foci of viral infection was the highest in NahG plants followed by RBOHD silenced plants, whereas lowest number of viral foci appeared in non-transgenic plants (Figure 5). These results could imply stronger impact of SA in immune signaling compared to RBOHD, one however has to note that NahG reduces the amounts of SA to less than 10% of native values (data not shown) while *RBOHD* is, in our transgenic lines, silenced to approximately 50% of its native transcriptional activity (Figure 5).

Although the oxidative burst following pathogen recognition occurs in the apoplast, being generated mostly by RBOH proteins and cell wall peroxidases, pathogen-induced ROS can be also produced in other cellular compartments like mitochondria, chloroplasts and peroxisomes (reviewed in Sharma et al., 2012; Mignolet-Spruyt et al., 2016). Our comparative analysis of NT and NahG genotypes shows that the extensive production of ROS corresponds to the spread of the virus and/or PCD (Figure 4; Lukan et al., 2018). Extensive ROS response in later stages of infection is most likely of non-apoplastic origin, as ROS production did not correspond to the expression of *RBOH* genes in NahG plants (Figure 5, Figure 6). This is also supported by the expression pattern of ROS quenchers CAT1 and PRX28, which are induced more distantly from the virus foci in NahG compared to NT plants (Figure 3). As SA was shown to be required for generation of mitochondrial ROS during pathogen infection (Liao et al., 2015), chloroplasts are its most probable source in our system. Transcriptional down-regulation of photosynthesis genes leads to generation of chloroplast ROS, which were proposed as a signal orchestrating PCD (Zurbriggen et al., 2010; Su et al., 2018). In our pathosystem transcriptional down-regulation of photosynthesis genes was observed (Baebler et al. 2014) thus we can assume that, similarly, chloroplastic ROS is generated. Our results of H_2_O_2_ staining (Figure 4) are in agreement with the hypothesis that chloroplastic ROS stimulates localised cell death.

ROS were also proposed to be a central component of a self-amplifying loop that regulates the interaction balance between different phytohormones such as SA, JA and ET (reviewed in Torres, 2010). SA regulates RBOHD-dependent ROS production in Arabidopsis (reviewed in Liu and He, 2016). The same is true for our pathosystem as induction of *RBOHD* expression was in NahG plants significantly reduced after PVY inoculation (Figure 5, Figure S6, Data S1). In Arabidopsis, RBOHD knockout mutant plants accumulated higher levels of SA following interaction with pathogen (Pogany et al., 2009). On the other hand, Chaouch et al. (2012) did not detect any difference in SA accumulation in RBOHD mutant compared to wild type Arabidopsis (Chaouch et al. 2012). We showed that in potato-PVY interaction, SA biosynthesis is not controlled by RBOHD generated ROS as the concentration of SA in transgenic plants with silenced RBOHD did not differ significantly from SA concentration in NT plants at the site of viral foci. RBOHD is however involved in spatial control of its accumulation (Figure 6) confirming the existence of complex regulatory feedback loops. Indeed, *UDP-glucosyltransferase* (*UGT76D1*) was identified as a component of the feedback activation loop of SA biosynthesis (Huang et al., 2018) and our results show that it is spatially repressed by RBOHD signalling, explaining the regulation of SA accumulation (Figure 6).

For efficient immunity plant needs to block the pathogen multiplication and its spread, and at the same time limit the extent of damaged tissue and energy consumption. Thus tight spatial regulation of this processes is of utmost importance. It has been long assumed that positive regulators act at the HR site and negative regulators in the surrounding areas, but the molecular evidence for this premise is mostly lacking due to lack of functional zonation studies (Huysmans et al., 2017). We here show that RBOHD and SA regulate the immune response in a spatio-diverse manner, leading to fine-tuning of both spatial and temporal response.

## Experimental Procedures

### Plant material

Potato (*Solanum tuberosum* L.) plants cv. Rywal and NahG-Rywal (Baebler et al. 2014) and derived transgenic lines (see below) were grown and inoculated with PVY^NTN^ (isolate NIB-NTN, AJ585342), PVY^N-Wilga^ (PVYN–Wi; EF558545) or with PVY^N605^, tagged with green fluorescent protein (PVY^N^-GFP) (Jakab et al., 1997; Dietrich and Maiss, 2003; Rupar et al. 2015) or mock-inoculated as described by Baebler et al., 2009.

### qPCR gene expression analysis in tissue sections

For gene expression analyses lesions at two different stages were sampled (see Figure 1a). For each lesion, 4 tissue sections were sampled (Figure 1a). Tissue sections were stored in 100 μl of RNAlater RNA Stabilization Solution (Thermo Fisher Scientific). Altogether 5 independent experiments were performed), each comprising of ca. 20 plants per experimental group. For each experimental group, 5-10 lesions from different plants were sampled (Table S3).

Standard gene expression analysis procedures were optimized for the analysis of small leaf sections. Fixed tissue sections were homogenized using Tissue Lyser (Qiagen), followed by RNA isolation using RNeasy Plant Micro Kit (Qiagen), DNAse treatment, quality control and reverse transcription.

Expression of 23 genes involved in different steps of immune signaling (Figure 1c) was analyzed and normalized to the expression of two validated reference genes *COX* and *18S* as described previously (Petek et al., 2014; see Table S4 for primer and probe information and full experimental details). The standard curve method was used for relative gene expression quantification using quantGenius (http://quantgenius.nib.si; Baebler et al., 2017).

### Statistical modelling of tissue section qPCR gene expression data

Prior to statistical analysis of tissue sections gene expression dataset, data from lesions without viral amplification detected in section A (necrosis due to mechanical inoculation) was filtered out. Relative gene expression values were next standardized to 97^th^ percentile. Multiple linear regression method, as implemented in R Stats package (R Core Team, 2017), was used to fit a quadratic polynomial model, with genotype as factor and position (tissue section) as a factor or numeric distance from the position A (initial viral foci tissue surrounding). Analysis of variance was additionally used to analyze the effects of position and genotype on gene expression. For each analyzed gene, spatial expression profiles were determined and plotted with 95% confidence intervals. For early visible lesions data, calculations were done for both perpendicular directions and their average, respectively. Expression changes between genotypes in different time post inoculation were further visualized using coefficient plots with linear coefficients displayed on the x-axis and quadratic coefficients on the y-axis. Significance of comparisons between the levels of the factors was obtained through contrast method as implemented in R package limma.

### Gene expression in whole leaves

For the gene expression analysis in whole leaves of different genotypes, including newly constructed transgenic lines (see experimental setups below), whole leaves were sampled. RNA was isolated, reverse transcription and qPCR was performed as described previously (Baebler et al. 2014), with normalization to COX only. Where applicable, t-test statistics (Excel) was used to compare treatments.

### RNA-Seq analysis in tissue sections

For RNA-seq, early visible lesions were sampled from PVY^N-Wilga^-inoculated leaves of cv. Rywal, NahG-Rywal and shRBOHD (line 14) plants. Thirty tissue sections of the lesion (A) and its immediate surrounding (B; Figure 1a) were collected and pooled separately (pool A, pool B). Sections of mock-inoculated plants were prepared as described above and pooled (cca 300 sections per pool). The pools were stored and homogenized as described above. RNA extraction and library preparation are described in Method S1. RNA-seq was performed on the HiSeq platform (Illumina) using 150-bp paired-end reads at Novogene or LC Sciences. Quality control was performed in CLC Genomics Workbench 12.0 (Qiagen). Merging of overlapping pairs, mapping the reads to the potato genome (Petek et al., 2019) and read counting were performed in both CLC Genomics and STAR. Differential expression analysis was performed in R (R Core Team, 2013; version 3.2.2), using the R package limma (Ritchie et al., 2015), TMM normalization and voom function. Adjusted p-values below 0.05 were considered statistically significant. See Method S1 for full experimental details. Raw and normalized RNA-seq data were deposited to GEO (accession number GSE142002, https://www.ncbi.nlm.nih.gov/geo/query/acc.cgi?acc=GSE142002; use password »axmbquqgvdgtpmp« for review).

Gene set enrichment analysis (GSEA; Subramanian et al., 2005) was performed to search for groups of genes involved in the same processes that were significantly (FDR corrected Q-value < 0.05) altered by virus inoculation, using MapMan ontology as the source of the gene sets (obtained from GoMapMan database, Ramšak 2014).

### Construction of short hairpin RNA transgenics

Primers were designed based on consensus sequence obtained by sequence alignment of sequences of *RBOHD* genes (Sotub06g025550 and Sotub06g025580) (Figure S10). cDNA obtained from cv. Rywal was used as a template for the amplification of the 436 bp-long fragment of *RBOHD* gene. The product was cloned into pENTR D-TOPO plasmid and sequenced. The fragment was further transferred to pH7GWIWG2, sequenced and electroporated into *Agrobacterium tumefaciens* LBA4404 that was used for transformation of cv. Rywal plantlets (see Figure S10 legend for full experimental details). Silencing efficacy was analyzed in one leaf per transgenic line by qPCR using shRBOHD primers (Table S4) as a target as described above.

### Phenotypisation

Lesions that developed on virus inoculated leaves of cv. Rywal, NahG-Rywal and two short hairpin transgenic lines with silenced RBOHD (lines 13 and 14) inoculated with PVY^N^-GFP or mock-inoculated were counted from 3 dpi to 14 dpi (see Data S5 for numbers of tested plants for each experiment). Numbers of lesions was calculated per cm^2^ of leaf area. For the same plants, times of lesion appearance on non-inoculated leaves was recorded.

Viral amount was analysed in same genotypes as above 6 dpi in the inoculated and 24 and 39 dpi in non-inoculated leaves (1-3 leaves from up to 5 plants). Sample preparation and qPCR analysis were performed as described above using PVY as a target gene. The experiment was repeated three times.

Lesion diameter was measured in the images of individual leaves of cv. Rywal and NahG-Rywal, inoculated with PVY^N-Wilga^ or PVY^NTN^ in Adobe Photoshop CS3 and standardized to leaf diameter. The experiment was repeated twice.

### H_2_O_2_ staining

The inoculated leaves (three leaves from three plants per genotype/treatment) of Rywal and NahG-Rywal plants, inoculated with PVY^NTN^ or mock inoculateds were sampled 3 to 8 days after inoculation and stained with DAB staining solution (see Method S2 for full experimental details). Diameters of lesions and DAB staining areas were measured on spatially calibrated images in Fiji (Schindelin et al., 2012). The difference in diameter of lesion and DAB area was measured in each individual lesion. Data was analyzed in R version 3.5.0 (R Core Team, 2018), statistical significance was tested with ANOVA and Tukey’s post-hoc test (p = 0.05).

### Hormonal measurements

SA and JA content were determined in non-transgenic cv. Rywal and two short hairpin transgenic lines with silenced *RBOHD* (lines 13 and 14) 4 dpi after PVY^N-Wilga^ or mock- inoculation. 60-130 sections containing initial virus foci (section A) and surrounding tissue (section B) were sampled from the third inoculated leaf from 6-8 plants of each genotype. Hormones were isolated and measured by gas chromatography coupled with mass spectrometry (GC-MS) as described by Križnik et al., (2017).

## Acknowledgements

The research was financially supported by the Slovenian Research Agency (research core funding No. P4-0165 and projects J4-7636, J4-1777 and 1000-15-0105) and Grant no. 2017/27/B/N29/01803 from NSC, Poland. The authors thank Dr. John Carr for critical review of the manuscript and Maja Gnezda, Neža Turnšek and Lidija Matičič for technical assistance.

## Author Contributions

ŠB, MPN, KG, TL, JH designed the research; ŠB, MPN, KG, TL, MTŽ, AK, KS, BD, AC, SP, KM, MK performed research; TL, ŠB, AK, MTŽ, AB, MZ, KG, MK, SP analyzed data; TL, ŠB, AK, MZ, KG, MK, MPN, MTŽ, BD, AC, JH contributed to the writing or the revision of the manuscript.

## Supplemental data files

- Figure S1: Symptoms appear systemically if Ny-gene-conferred resistance is perturbed in SA signaling.
- Figure S2: Lesion diameter is two-fold larger in NahG compared to NT plants following inoculation with PVY^N-Wilga^ or PVY^NTN^.
- Figure S3: Transcriptional response of selected immune signaling-related genes and relative viral RNA abundance is similar after inoculation with different viral strains.
- Figure S4: Consistent spatial gene expression of immune signaling genes around early visible lesions.
- Figure S5: 9-LOX and 13-LOX branches of the oxylipin pathway respond differentialy in potato resistance against PVY.
- Figure S6: Virus is present in symptomatic non-inoculated leaves of plants of RBOHD-silenced transgenic lines.
- Figure S7: Lesions on inoculated leaves of plants of different genotypes.
- Figure S8: Spatial transcriptional response is similar in shRBOHD and NahG genotypes.
- Figure S9: Spatial expression of UDP-glucosyltransferase (UGT76D1) in different genotypes.
- Figure S10: Construction of short hairpin RNA transgenics.
- Table S1: Gene expression changes of selected genes in whole leaves.
- Table S2: Sample information and salicylic acid content in tissue sections of NT and two shRBOHD lines.
- Table S3: Sampling for spatiotemporal gene expression analyses.
- Table S4: Selected genes and corresponding primers and probes sequences for expression analyses with quantitative PCR.
- Data S1: Sample information and gene expression of selected genes in lesion sections.
- Data S2: Statistical modeling of tissue section gene expression data.
- Data S3: Spatiotemporal gene expression of immune signaling genes.
- Data S4: 3D animated spatial gene expression of immune signaling genes around early visible lesions.
- Data S5: Number of lesions on inoculated leaves of different potato genotypes following PVY inoculation.
- Data S6: Gene expression changes of Rywal, NahG and shRBOHD genotypes following PVY^N-Wilga^ infection determined by RNA-Seq.
- Methods S1: RNA-Seq analysis.
- Methods S2: H_2_O_2_ staining and imaging.

